# Elevated temperatures impose transcriptional constraints on coffee genotypes and elicit intraspecific differences in thermoregulation

**DOI:** 10.1101/2020.03.07.981340

**Authors:** Raphael Ricon de Oliveira, Thales Henrique Cherubino Ribeiro, Carlos Henrique Cardon, Lauren Fedenia, Vinicius Andrade Maia, Barbara Castanheira Ferrara Barbosa, Cecílio Frois Caldeira, Patricia E. Klein, Antonio Chalfun-Junior

**Author notes:** These authors contributed equally to this study (Raphael Ricon de Oliveira, Thales Henrique Cherubino Ribeiro).

## Abstract

The projected impact of global warming on coffee production may require the heat-adapted genotypes in the next decades. To identify thermotolerance cellular strategies, we compared the effect of elevated temperature on two commercial *Coffea arabica* L. genotypes exploring leaf physiology, transcriptome and carbohydrate/protein composition. Growth temperatures were 23/19°C (day/night), as optimal condition (OpT), and 30/26°C (day/night) as a possible warmer scenario (WaT). The cv. Acauã showed lower levels of leaf temperature under both conditions compared to cv. Catuaí, whereas slightly or no differences for other leaf physiological parameters. Therefore, to explore thermoregulatory pathways the leaf transcriptome was examined using RNAseq. Genotypes showed a marked number of differentially-expressed genes (DEGs) under OpT, however DEGs strongly decrease in both at WaT condition indicating a transcriptional constraint. DEGs responsive to WaT revealed shared and genotype-specific genes mostly related to carbohydrate metabolism. Under OpT, leaf starch content was greater in cv. Acauã although the levels of leaf starch, sucrose, and leaf protein decreased in both genotypes as WaT was imposed. These findings indicate that genotypes with a greater capacity to maintain carbohydrate homeostasis under temperature fluctuations could be more thermotolerant and which may be useful in breeding for a changing climate.

**HIGHLIGHT:** In response to warming, transcriptional differences decrease in coffee genotypes hampering breeding programs. Differences in gene expression and sugar levels confirm intraspecific variation associating thermotolerance to maintenance of energetic homeostasis.

## INTRODUCTION

Climate change is multifaceted and despite measurable impacts of elevated temperatures on agriculture (Lobell *et al*., 2011; Zhao *et al*. 2017), there remains considerable gaps on how coffee systems will be affected by both short- and long-term changes in the environment. Several studies on the impact of climate change on coffee systems have projected marked negative effects on yield, berry quality, suitable planting areas, and incidence of disease and insects. Collectively, these environmental stresses will likely impose both economic and social problems within many coffee producing regions (reviewed by DaMatta *et al*., 2019). Despite attenuating factors associated with increasing global CO_2_ levels that could partially mitigate the negative production trends described above (Rodrigues *et al*., 2016) and numerous studies demonstrating the impact of temperature on coffee physiology (Drinnan and Menzel, 1995; DaMatta and Ramalho, 2006; Läderach *et al*., 2017), a detailed understanding of the molecular thermoregulatory mechanisms is lacking.

*C. arabica* L. is a tropical tree responsible for the major worldwide production of coffee (ICO, 2019) and its optimal growth temperature is considered between 18 to 23°C (Camargo, 1985; Teketay, 1999). The coffee tree has a periodicity growth habit that closely follows rainfall patterns and, historically, it is considered highly sensitive to climatic changes, especially temperature and drought (DaMatta and Ramalho, 2006; Camargo, 2010; DaMatta, 2018). Mean temperatures are projected to increase by 2.6 - 4.8 °C (IPCC, 2013; IPCC, 2014), which may have serious repercussions on coffee production. Considering these changing temperatures, select genotypes were identified that outperformed others when exposed to higher annual mean temperatures. This suggests there is potentially useful intraspecific variability of thermotolerance in some genotypes and investigation into the molecular mechanisms underlying this variability is warranted (DaMatta, 2018).

Increasing temperature impacts plant physiology from the cellular to the whole plant level and changes photoassimilate allocation to repair and recovery processes (Bita and Gerats, 2013; Bokszczanin *et al*., 2013; Marias *et al*., 2017a). However, the stress severity depends on intensity and duration of exposure beyond the plant species and within genotypes (Teskey *et al*., 2015). Therefore, beyond particular characteristics and growth conditions, a comprehensive effect of increasing temperature on plants needs first to differentiate data from a moderate long-term change to more drastic ones such as short-duration heat-waves (Thornton *et al*., 2014). Both phenomena are predicted to be more frequent in the future and may occur singly or concomitantly (Hao *et al*., 2013; IPCC, 2014) highlighting the need for independent and overlapping studies.

To understand the limits of coffee thermotolerance, recent studies have explored the effect of a gradual increasing temperature or extreme heat stress on select physiological processes (reviewed by DaMatta *et al*., 2019). Minimal impact on photosynthetic-related parameters was observed when various coffee genotypes were exposed to temperatures up to 37°C whereas maximum photosynthetic damage occurred at 42°C for all coffee genotypes (Martins *et al*., 2016). Although coffee presents moderate thermotolerance of photosynthetic-related processes, most genotypes produced abnormal reproductive structures at these elevated temperatures (DaMatta *et al*., 2019). Accordingly, coffee plants subjected to 45°C for 1-to-1.5 hours showed leaf age-related differences in physiological recovery and did not bear flowers or fruits (Marias *et al*., 2017a). These results demonstrate that, depending on the tissue and stage of plant development, coffee thermotolerance may be substantial regarding physiological parameters.

From the molecular point of view, temperature is perceived by multiple pathways in model plants and crops (Wigge, 2013; Hasanuzzaman *et al*., 2013; Jagadish *et al*., 2016; Ibañez *et al*., 2017). A general and critical cellular response to heat stress is the activation of heat shock proteins (HSPs), which function as chaperones ensuring proper folding of proteins (Ohama *et al*., 2016). Importantly, phytochromes act as thermosensors joining the related processes of light perception to temperature (Jung *et al*., 2016).

The impact of elevated temperature on gene expression and associated thermotolerance is highly heterogeneous in plant species (von Koskull-Döring *et al*,, 2007; Ohama *et al*., 2017). Therefore, extrapolation of molecular mechanisms relating to thermotolerance in model species is unreliable and will require direct validation. At present, molecular studies examining the effect of elevated temperature on *Coffea* sp. are limited when compared to molecular-based drought studies in this crop (Moffato *et al*., 2016; DaMatta *et al*., 2019). To the best of our knowledge, a large-scale analysis of the intraspecific transcriptional variation in response to elevated temperature has not been reported in *Coffee arabica* L. and could reveal thermotolerance molecular pathways and important strategies towards breeding programs.

Thermotolerance is acquired via protective cellular machinery gained throughout coffee plant maturation (Marias *et al*., 2017a; 2017b). This suggests that young plants are more sensitive and require long-term exposure to stress to acclimatize making this stage in plant development a useful model to examine the impact of warmer temperatures on gene expression during acclimation. Thus, to study intraspecific variation associated with mechanisms of thermotolerance on coffee, the present study imposed elevated temperatures on 1-year old plants of two coffee genotypes, cv. Catuaí IAC 144 and cv. Acauã, which have been reported to differ in thermotolerance (Carvalho, 2008). Physiological parameters were evaluated as well as a global transcriptional analysis in conjunction with an initial metabolomics investigation of photo-assimilates, sugars and protein.

## MATERIAL AND METHODS

### Plant material

Two *Coffea arabica* genotypes, cvs. Acauã and Catuaí IAC 144, hypothesized to differ in heat tolerance (Carvalho, 2008) were examined in the present study. Coffee plants were cultivated in growth chambers with 12 hours of light, 60% humidity and either 23/19°C or 30/26°C (day/night temperatures) that are considered the optimal (OpT) and warm temperatures (WaT), respectively (DaMatta and Ramalho, 2006). For the RNAseq and RT-qPCR analyses, plants were obtained from two hundred seeds of each genotype cultivated for thirty days in a commercial substrate (Professional Growing Mix, Sun-Gro Horticulture). After thirty days, seedlings were individually transplanted to a two-liter-pot and maintained in greenhouses (Department of Horticultural Sciences, Texas A&M University, USA) with 50% shade until the three leaf pair stage. At the three paired leaf stage, plants were transferred to the Texas A&M AgriLife Research and Extension Center (Overton, TX), randomized in complete blocks with split plot restrictions. Plants were allowed to acclimate under control growth conditions for fifteen days at OpT. Acclimated coffee plants were then divided between two chambers at either OpT or WaT conditions and maintained for four weeks. Each biological repetition was comprised of five excised leaves that were harvested immediately and placed in liquid nitrogen, pulverized with a mortar and pestle, and subsequently stored at -80 °C until analysis.

For gas exchange and sugar content analyses, the experiment was repeated using similar age plants of each genotype transferred from greenhouses to a Conviron^®^ growth chamber (Plant Physiology Sector, Federal University of Lavras, Brazil). Twenty plants were transplanted to a mix of soil, sand and fertilizer formula 5–25-15 of N-P-K and maintained in a greenhouse for one week. Then, they were transferred to a chamber, acclimatized for two weeks at OpT with 12 hours of light and maintained for four weeks at OpT and after four weeks at WaT.

### Gas exchange measurements

Physiological parameters, such as carbon assimilation rate, stomatal conductance, transpiration rate and chlorophyll fluorescence were measured for each coffee genotype at different temperatures and maintained between the second and fourth hours of the light period on completely expanded leaves. Ten plants of each cultivar were randomly selected, and one leaf from each used for measurements with a portable infrared gas analyzer IRGA (LI-6400XT, LI-COR^®^) once a week for four weeks. These measurements were done with reference CO_2_ concentration fixed at 400 µM mol_-1_, relative humidity was set to 60% and photon flux density inside the measuring chamber to 1000 µmol m_-2_s_-1_.

### RNAseq library preparation

Five biological repetitions for each genotype at the two growth temperature regimes were used (20 RNAseq libraries). The RNA extractions were performed with 100 mg of powdered tissue using the Concert^™^ Kit Plant RNA Reagent (Invitrogen^®^) and followed by treatment with the Turbo DNA-free Kit (Ambion^®^). RNA integrity and purity were assessed by 1% agarose gel electrophoresis and analyzed on a DeNovix DS-11 spectrophotometer (DeNovix Inc., Wilmington, DE, USA) and a Bioanalyzer 2100 (Agilent Technologies, Santa Clara, CA). All samples presented standard values and RNA Integrity Number (RIN) higher than 7.0. The TruSeq library preparations were constructed using the cDNA Synthesis kit (Illumina Inc., San Diego, CA, USA). Two lanes of paired-end (2×150 bp) sequencing of the cDNA libraries were performed on the Illumina HiSeq 2000 (Illumina Inc., San Diego, CA, USA). Library preparation and sequencing were performed by AgriLife Genomics and Bioinformatics Services (Texas A&M University, College Station, TX, USA) in April 2015. Sequence cluster identification, quality prefiltering, base calling and uncertainty assessment were done in real-time using Illumina’s HCS 2.2.58 and RTA 1.18.64 software with default parameter settings. All the reactions followed the respective manufacturer’s instructions. Pre-processed libraries are available in SRA under BioProject ID PRJNA609253.

### RNAseq analysis

Approximately 183 million sequenced paired-end reads were used for alignment against the *Coffea canephora* genome (available at http://coffee-genome.org) using the STAR v. 2.5.3a aligner with default parameters. Libraries were sorted and PCR duplicates were removed with Picard tools. Approximately 115 million paired-end reads were uniquely mapped to exons and read counts were quantified with htseq-count script. For differential expression analyses the library WAT_AC_1 (cv. Acauã at WaT conditions, replicate 1) was not considered due to a relative low number of uniquely mapped reads (∼2.3 million). Differentially expressed genes (DEGs) of the same cultivar in contrasting temperature conditions were identified using the Bioconductor R package *edgeR* by comparing the normalized number of reads aligned to each gene model in different conditions using a Generalized Linear Model applied to the expression matrix (Robinson *et al*., 2010; Huber *et al*., 2015). Benjamini and Hochberg’s false discovery rate (FDR) below 0.05 and a minimum log_2_ fold change of one were the parameters used to consider a gene differentially expressed between the two conditions. To improve the quality of functional characterization of the DEGs, their respective protein sequences were subjected to homology searches with BLASTP version 2.7.1+ (Camacho *et al*., 2009) against all plant proteins in the NCBI non-redundant protein database (nr). In addition, we enriched our DEG results by mapping those proteins against the KEGG database with the BlastKOALA tool (Kanehisa *et al*., 2016) in order to find the pathways that the DEGs were related to.

### Quantitative gene expression analysis (RT-qPCR)

RT-qPCR analysis was conducted from three biological repetitions with two technical replicates for each genotype at both growth temperatures. Total RNA was isolated using 100 mg of frozen powdered tissue and the PureLink™ Plant RNA Reagent System (Thermo Fisher^®^, Invitrogen) according to the manufacturer’s protocol. Samples were subsequently treated with the Turbo DNA-free Kit (Ambion^®^) for removal of DNA contamination. RNA purity was analyzed on a DeNovix DS-11 spectrophotometer (DeNovix Inc.). First-strand cDNA was synthesized using SuperScript® III First-Strand Synthesis System (Invitrogen™) according to the manufacturer’s protocol. RT-qPCR reactions were conducted using SYBR Green MasterMix (Applied Biosystems^®^) following the manufacturer’s instructions. Gene-specific primers (Table S2) were designed in non-conserved regions using the Primer-BLAST tool (Ye *et al*., 2012) with primer specificity validated using the CoffeeHub and Phytozome databases (Goodstein *et al*., 2012; Denoeud *et al*. 2014). Primer efficiency and RT-qPCR analyses were performed using the CFX384 Touch™ Real-Time PCR Detection System (Bio-Rad Laboratories, Hercules, CA, USA). Differential gene expression analysis was inferred using an adapted modeling approach under dCt values (Yuan *et al*., 2006) in relation to the reference genes *Malate dehydrogenase* (*MDH*, GW464198.1) and *Ubiquitin-conjugating enzyme E2* (*UBQ2*; GR984245) previously described and validated for RT-qPCR in *Coffea* spp. (Martins *et al*., 2017).

### Carbohydrate and protein content

Carbohydrate and protein content analyses were conducted from four biological repetitions with two technical replicates for each genotype at the two growth temperatures. The extraction of carbohydrates and proteins was based on Zanandrea *et al*. (2010) with modifications (Silva *et al*., 2014) in which 1000 mg of frozen powdered tissue (fresh weight) were homogenized in 5 mL of 100 mM potassium phosphate buffer (pH 7.0) and then placed in a water bath for 30 min at 40 °C. The solution was centrifuged at 10,000 g for 10 min and the supernatant was collected. The process was repeated twice and supernatants were combined totalyzing 10 ml. For extraction of starch, the pellet was resuspended in 10 mL of 200 mM potassium acetate buffer (pH 4.8) and 16 units of amyloglucosidase enzyme were added. Then, samples were incubated in a water bath at 40 °C for two hours. Following centrifugation at 10,000 g for 20 min, the supernatant was collected for measurements. Starch, sucrose and total soluble sugars were quantified as described by Dische (1962), and the level of reducing sugars was quantified according to Miller (1959). Protein was quantified as described by Bradford (1976) and analyzed in a spectrophotometer at 570 nm comparing results with a standard curve of 0.1 μmol/mL Bovine Serum Albumin (BSA).

### Statistical analysis of Physiologic, Metabolic and RT-qPCR expression data

The modeling approach was carried out by Linear Mixed Models (LMM) using the “lmer” function from the lme4 R package (Bates *et al*., 2015) for the statistical analysis of IRGA physiological data, metabolic parameters and RT-qPCR expression. In all experiments individuals were used as random factors to deal with the dependence between observations at the same individual across different weeks or conditions. Additionally, the models were fitted by maximum likelihood. The treatments were coded as a factor level and used as fixed effects including temperature conditions (WaT or OpT), cultivar (Catuaí or Acauã) and weeks (only for the physiological analyses), in cases of interest, the interactions between the fixed effects were accounted. Residuals normality and variance homogeneity were assessed by Shapiro-Wilk test and residuals versus fitted plots, respectively. The post hoc pairwise contrasts between factor levels were obtained by “lsmeans” function from “lsmeans” package (Lenth, 2016) using Tukey adjust method. Statistical significance was assessed using Satterthwaite approximation, by the package lmerTest (Kuznetsova *et al*., 2015). For the RT-qPCR analysis, genes were considered differentially expressed if their expression profile respected three parameters: 1) the residuals of modeled Ct values presented a normal distribution, 2) pairwise differences of two contrast conditions (i.e. WaT Acauã plants against OpT Acauã plants) presented a Tukey adjusted p-value<0.05, and 3) the expression mean of a given gene was at least 2 times more expressed in one of the compared conditions (−1 < log_2_FC > 1).

## RESULTS

### Coffee genotypes present thermoregulatory differences in response to warm temperatures

Physiological analyses showed that coffee genotypes, cv. Catuaí and cv. Acauã, present similar trends under optimal temperatures (OpT) and also in response to warm temperatures (WaT), however quantitative and transient differences were observed for the two genotypes during the 4 week experiment (Fig. 1 and Table S1 for statistical analyses). The main quantitative difference was noted for leaf temperature (Tleaf) where cv. Catuaí consistently had higher temperature values than cv. Acauã independent of the imposed conditions (Fig. 1A).

**Fig. 1–.**
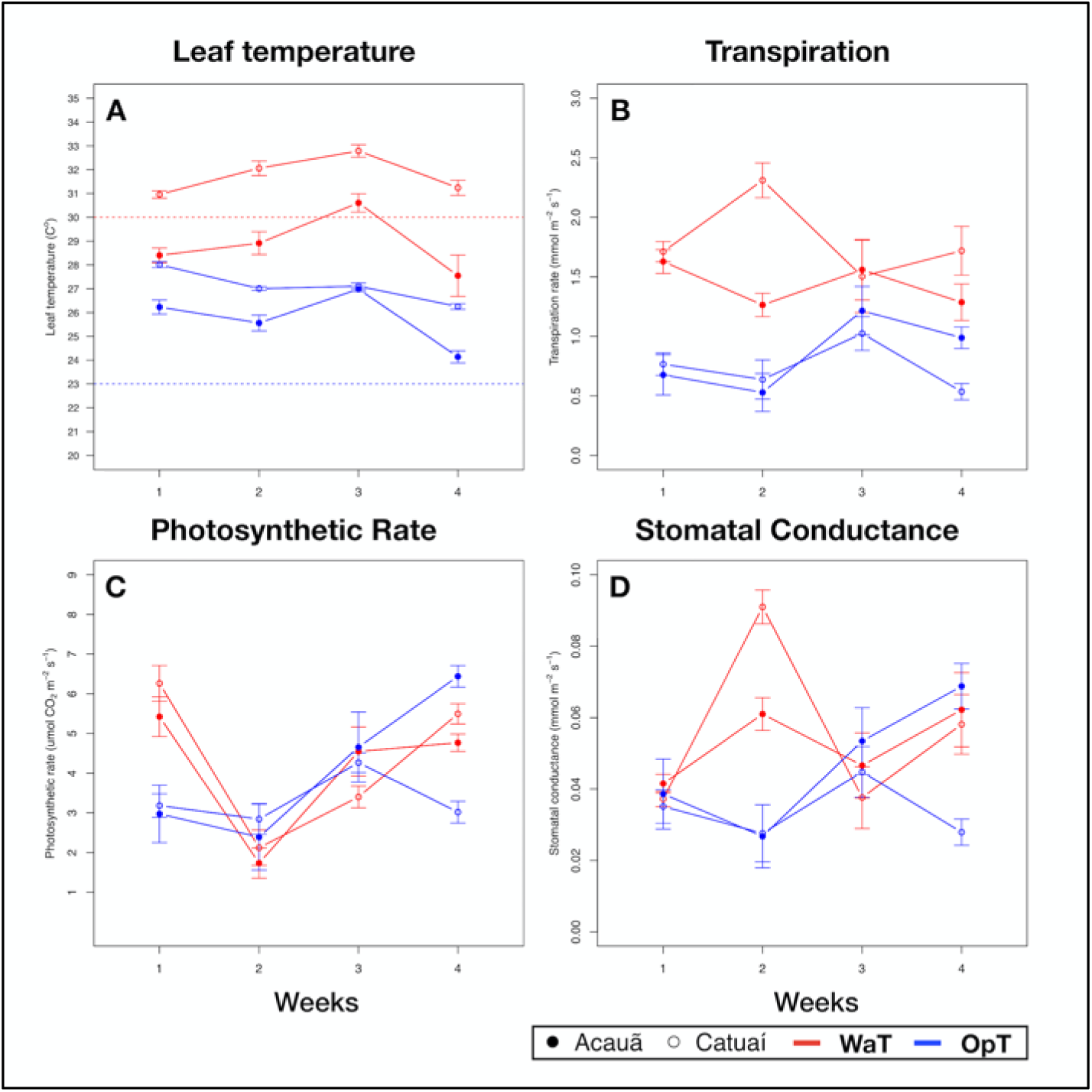
Physiological analysis of two coffee genotypes under optimal and warm temperatures. Physiological parameters of two coffee genotypes, cv. Acauã (closed circles) and cv. Catuaí (open circles), were measured along four weeks for each temperature condition, 23/19°C (OpT) and 30/26°C (WaT). A) Leaf temperature (Tleaf). Dotted lines in the figure represent the chamber ambient temperature, OpT (blue dotted line) or WaT (red dotted line), at the time measures were made; B) Transpiration rate; C) Photosynthetic rate; D) Stomatal conductance. Labels: OpT - optimal temperature (23/19°C, day/night); WaT - warm temperature (30/26°C, day/night).

At OpT conditions, both genotypes had Tleaf above 23 °C throughout the 4-week treatment, however, cv. Catuaí kept Tleaf between 26-28 °C whereas Acauã always had a lower Tleaf than cv. Catuaí, except at week 3 under OpT (Fig. 1A and Table S1). These differences in Tleaf were not correlated with leaf transpiration as both genotypes showed similar values at OpT (Fig. 1B). From this, we concluded that plants of cv. Catuaí, in general, presented a basal temperature higher than cv. Acauã. As plants were subjected to WaT conditions, Tleaf gradually increased for both coffee cultivars during the first three weeks of elevated temperatures and then decreased for both cultivars (Fig. 1A). As was observed under OpT conditions, cv. Catuaí showed higher Tleaf values compared to cv. Acauã. Catuaí always remained above the 30°C imposed by chamber ambient temperature whereas cv. Acauã showed Tleaf mostly below 30°C.

In examining leaf transpiration at WaT, both cultivars displayed an increase in transpiration over that observed at OpT (Fig. 1B). A transient difference in transpiration was observed at week 2 with cv. Catuaí showing a marked increase, but this difference was not significant and did not persist into the subsequent weeks under elevated temperature (Table S1). Regardless, Tleaf and transpiration values did not correlate well since, in general, lower leaf temperatures are associated with evaporative cooling driven by higher transpiration rates. This poor correlation is especially apparent when examining the results observed at week 2 under WaT conditions; transpiration rate and leaf temperature for cv. Catuaí was markedly higher compared to cv. Acauã (Fig. 1A, 1B and Table S1). We propose that cv. Catuaí juvenile trees have a lower thermoregulatory efficiency because Tleaf was higher than cv. Acauã under both OpT and WaT growth conditions.

Additionally, photosynthetic rates and stomatal conductance did not show consistent differences across time points between the coffee genotypes (Fig. 1C, 1D and Table S1). For instance, comparing the two genotypes in each condition a similar trend can be noted, except at week 4 for OpT conditions where a significant difference was observed (Fig 1C, Table S1). For stomatal conductance, consistent differences between the two genotypes were not apparent since stomatal conductance varied only at week 4 in OpT (Fig. 1D and Table S1). These results suggest that, despite apparent differences in their thermoregulatory efficiency, the coffee genotypes examined did not show consistent differences in photosynthesis or stomatal conductance during the 4 weeks of elevated growth temperature.

### Transcriptional pathways related to energy metabolism are affected by warmer temperatures in a genotype-dependent manner

The general response of plants to temperature stress involves multiple biological processes including transcriptional reprogramming and changes in cellular/physiological processes (Mittler *et al*., 2012; Barah *et al*., 2013). Since the present results indicate that coffee cv. Acauã and cv. Catuaí possess thermoregulatory differences in leaves (Fig. 1), we conducted a global transcriptome analyses to characterize the capacity of these coffee genotypes to respond to elevated temperatures through molecular regulatory pathways. We performed RNAseq analysis on leaf tissue from the two coffee cultivars and characterized differentially expressed genes (DEGs) within and between the two cultivars in response to warming (Fig. 2).

**Figure 2–.**
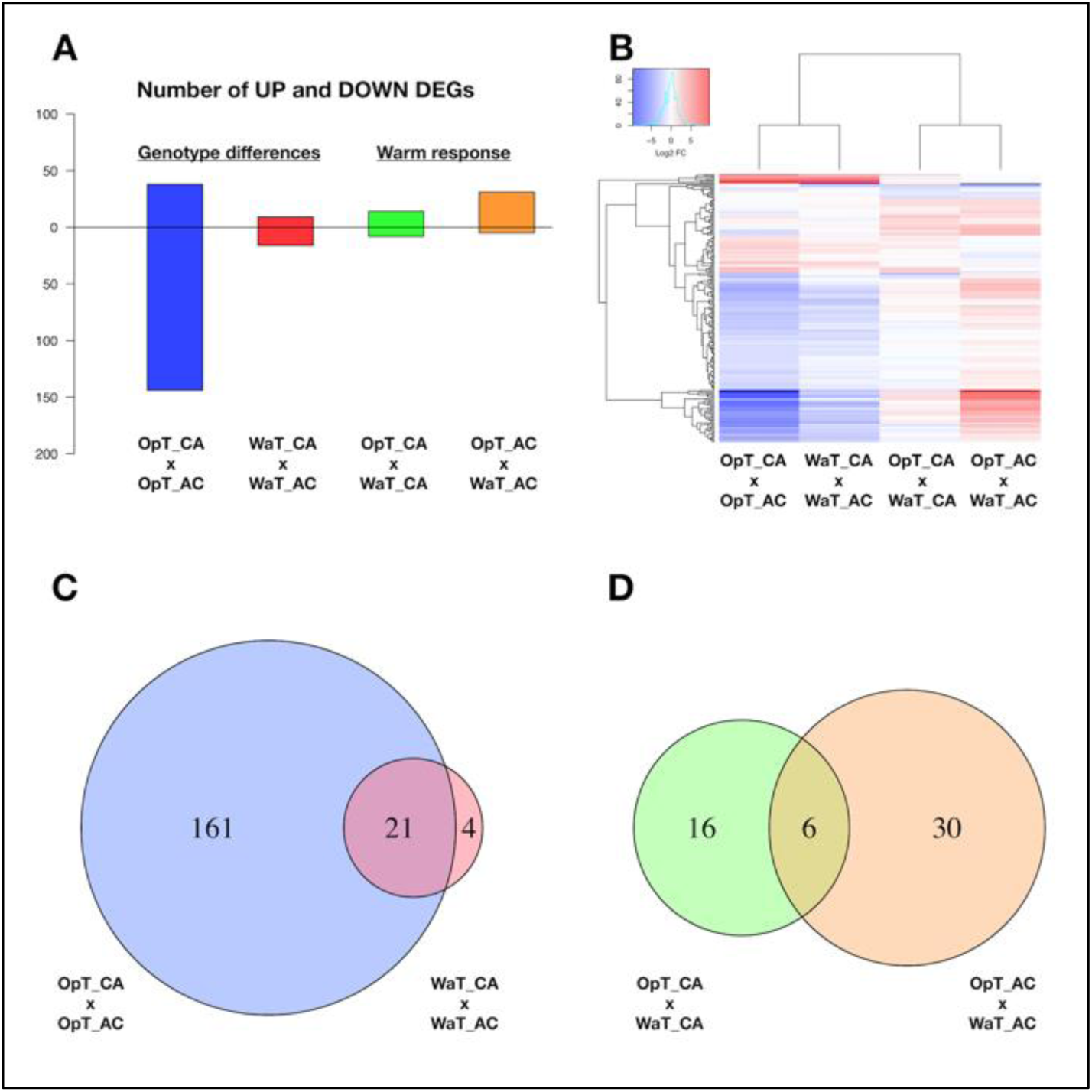
RNAseq analysis of two coffee genotypes under two temperature conditions. A) Number of differentially expressed genes (DEGs) between coffee genotypes, cvs. Acauã and Catuaí, at stated temperature (OpT or WaT; named “Genotype differences” in figure) and between the same cultivar at different temperatures (named “Warm response”). B) Heat map with up and down-regulated DEGs shown in A. C) Venn diagram showing the DEGs found between coffee genotypes at OpT and WaT conditions (Genotype differences) highlighting that the warm temperature causes a drastic reduction on DEGs. D) Venn diagram showing the DEGs responsive to WaT for each coffee genotype (Warm response). Details of DEGs, including genome identification, annotation and expression analysis, are available in Table S2. Labels: OpT - optimal temperature (23/19°C, day/night); WaT - warm temperature (30/26°C, day/night); CA - cv. Catuaí; AC - cv. Acauã.

Two types of RNAseq analyses were made. One analysis compared gene expression between the two coffee cultivars at a select temperature (OpT; WaT), which revealed DEGs related to transcriptional differences between genotypes at a given growth temperature (Fig. 2A). The second analysis examined DEGs within a genotype in response to different temperature conditions, which revealed genotypic-dependent DEGs in response to WaT (Fig. 2A). A heat map shows the expression levels of DEGs ranging between -10 to +10 fold changes in expression, and presents two visible patterns; most of the DEGs were down-regulated in cv. Acauã in relation to cv. Catuaí at a fixed growth temperature, whereas most of DEGs responsive to WaT were up-regulated in both genotypes (Fig. 2B). The annotation and fold expression details of all DEGs are provided in Table S2. In total, 186 DEGs were found when comparing gene expression of cv. Acauã to cv. Catuaí (Fig. 2C and Table S2); 161 DEGs were observed at OpT (130 down- and 31 up-regulated), 4 DEGs were detected exclusively at WaT (2 down- and 2 up-regulated), and 21 DEGs were shared across the two temperature regimes (14 down- and 7 up-regulated).

To functionally characterize the DEGs, we performed analysis of gene ontology (GO) and pathways using blast2GO (Götz *et al*., 2008), KEGG toolkits (Kanehisa *et al*., 2016) and AgriGO (Tian *et al*., 2017). The main categories found for DEGs were related to carbohydrate and protein metabolism (Fig. S1A), while biological process and molecular function of DEGs were related to biotic stimulus, defense response, oxi-reduction and oxidoreductase activity (Fig. S1B). In agreement, the pathway differences of starch and sugar metabolism (Fig. S2) showed DEGs related to enzymes including Sucrose-6-phosphate (EC 3.2.1.26), UDP-Glucose (EC 2.4.1.13) and Trehalose (EC 3.1.3.12). Thus, coffee genotypes at one-year of age presented differences in gene transcription at optimal growth temperatures that are related to energy metabolism. In contrast, the number of DEGs between the two genotypes were reduced drastically when cultivars were placed under WaT (Fig. 2C) suggesting that many of the intraspecific transcriptional differences were restricted to optimal temperature conditions.

Our analyses of coffee genotypes revealed a total of 52 DEGs in response to WaT (Fig. 2D and Table S2), in which 16 DEGs occurred exclusively in cv. Catuaí (9 up- and 7 down-regulated) and 30 in cv. Acauã (26 up- and 4 down-regulated) while 6 DEGs were in common between the two genotypes (5 up- and 1 down-regulated). We performed gene annotation and GO analyses of DEGs (Table S2 and Fig. S3) which revealed that the most enriched GO category and biological process was related to carbohydrate metabolism (Fig. S3). Indeed, three of these DEGs represent enzymes that are part of the carbohydrate pathway of starch and sucrose metabolism (Fig. S4); Granule-bound starch synthase (EC 2.4.1.242/Cc08_g16970), Glucose-1-phosphate adenyltransferase (EC 2.7.7.27/Cc02_17340) and α-amylase (EC 3.2.1.1/Cc06_g08480).

In cv. Acauã, we found additional molecular pathways represented by DEGs in response to warmer temperatures that included plant hormone signal transduction and carbon metabolism (Fig. S5 and S6), represented, respectively, by the DEGs ABA responsive element Binding Factor (ABF; Cc10_g04070) and Pyruvate Phosphate DiKinase (PPDK; Cc03_g02730; EC:2.7.9.1). These results are in agreement with the higher number of DEGs responsive to WaT found in cv. Acauã (Fig. 2D).

To compare expression differences between genotypes in response to WaT and possible implications on metabolic regulatory pathways, we selected 10 DEGs to check gene expression (out of 52) related to energy metabolism and thermotolerance, in which six were shared and three exclusive to cv. Acauã and one exclusive to cv. Catuaí (Fig. 2D). Nine of these 10 DEGs showed upregulation in at least one coffee genotype in response to WaT, whereas the *Small Heat shock* (*sHSP-like;* Cc11_g16360) was only downregulated in both. *PCC13-62* was the only DEG that presented expression difference between genotypes at the WaT conditions. The RNAseq results for these DEGs were validated by RT-qPCR, which showed similar expression trends for the 10 DEGs (Fig. S7, see Table S3 for statistics).

DEGs between coffee genotypes (Fig. 2D) suggest the existence of a conserved mechanism in response to WaT and also exclusive pathways, both mainly related to carbohydrate metabolism control (Fig. S1 and S3). For example, *Granule-bound starch synthase 1* (Cc08_g16970) was up-regulated in both coffee genotypes in response to WaT (Fig. 3 and S4). However, other regulatory genes such as *Glucose-1-phosphate adenyltransferase* (Cc02_17340) was only observed up-regulated in cv. Acauã whereas *Alpha-amylase* (Cc06_g08480) was only up-regulated in cv. Catuaí. These results demonstrated that the transcriptional pathways related to energy metabolism are affected by warmer temperatures in a genotype-dependent manner, consistent with results comparing different coffee species (Bertrand *et al*., 2015).

**Figure 3–.**
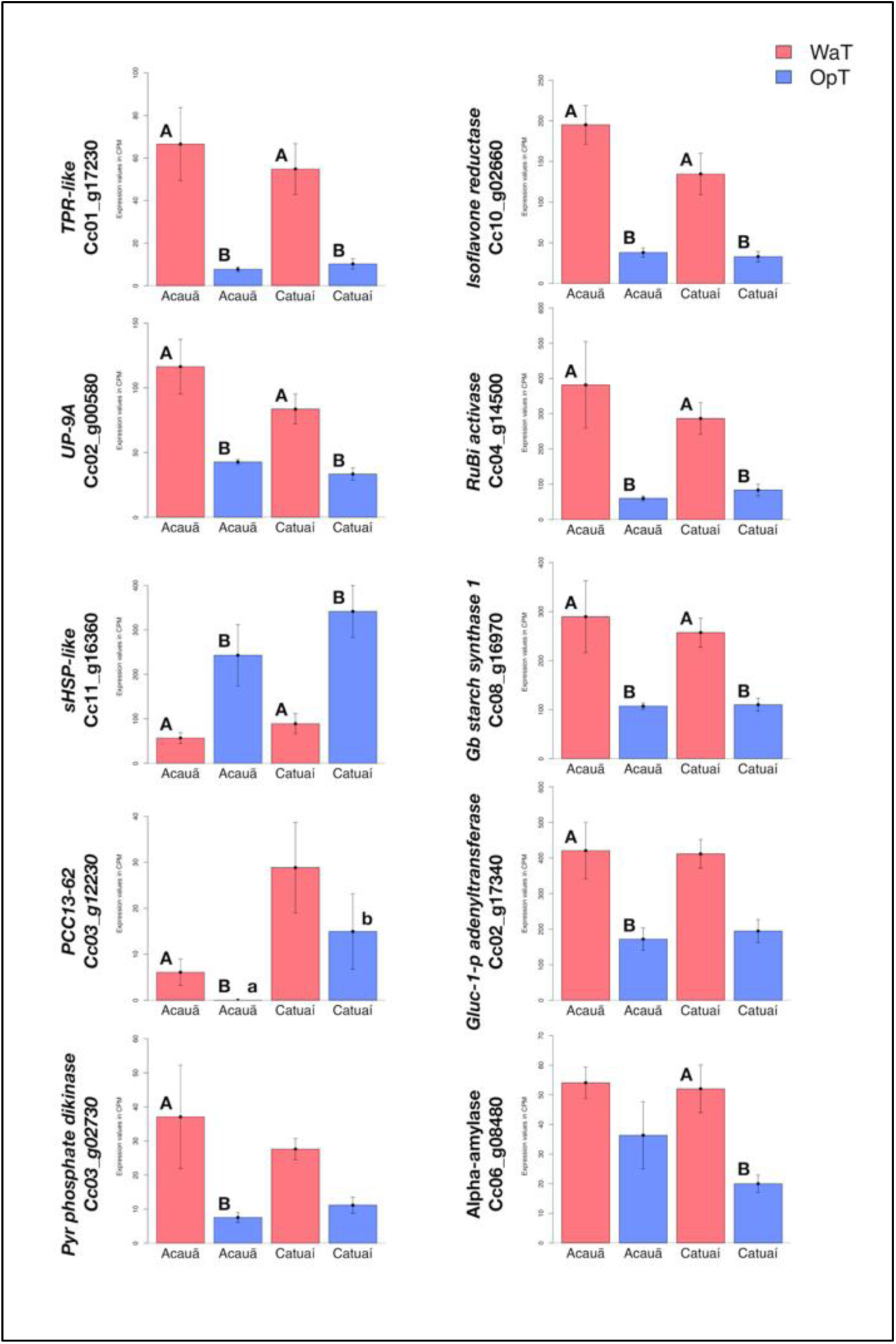
RNAseq expression analysis of warm-responsive DEGs related to energy metabolism and thermotolerance. Based on analysis of DEGs (Fig. 2D and Table S2), we evaluated RNAseq expression of ten DEGs related to energy metabolism and thermotolerance; six DEGs that were shared between the two genotypes and four that were exclusive to one of the two genotypes, three in cv. Acauã and one in cv. Catuaí, respectively: the *TPR-like* (Cc01_g17230), *Isoflavone reductase* (Cc10_g02660), *UP-9A* (Cc02_g00580), *RuBisCO activase* (*RuBi activase;* Cc04_g14500), *Small Heat Shock Protein like* (*sHSP-like;* Cc11_g16360) and *Granule-bound starch synthase 1* (*Gb starch synthase 1;* Cc08_g16970); *Desiccation-related_protein_PCC13-62* (*PCC13-62*; Cc03_g12230), *Glucose-1-phosphate adenyltransferase* (*Gluc_1_p_adenyltransferase*; Cc02_17340); *Pyruvate phosphate dikinase* (*PPDK*; Cc03_g02730); and *Alpha-amylase* (Cc06_g08480). Statistical analyses were performed comparing the same coffee genotype at different temperatures (capital letters) and comparing different genotypes at the same temperature (small letters). Differences were considered significant at p<0.05 (see Table S2 for details). Error bars represent standard errors. Labels: OpT (blue columns) - optimal temperature (23/19°C, day/night); WaT (red columns) - warm temperature (30/26°C, day/night).

### Warmer temperature affects sugar and protein content of coffee genotypes

Based on previous results from transcriptional analyses, we investigated whether warmer growth temperatures could affect the sugar content in a genotype-dependent manner (Fig. 4). With several noted exceptions, i.e. soluble and reducing sugars in cv. Acauã leaves, both coffee genotypes showed similar patterns with higher carbohydrate and protein content in leaves at OpT compared to WaT growth conditions (Fig. 4). Statistical analyses revealed specific differences in response to WaT (see Table S4), including a significant drop in leaf starch content in cv. Acauã at WaT (p<.0001; Fig. 4A), whereas starch content decreases in cv. Catuaí was significant at a much lower probability level (P<.0628). Accordingly, cv. Acauã showed higher leaf starch levels at OpT conditions than cv. Catuaí, but leaf starch content dropped to a similar low level in both cultivars when exposed to WaT conditions.

**Figure 4–.**
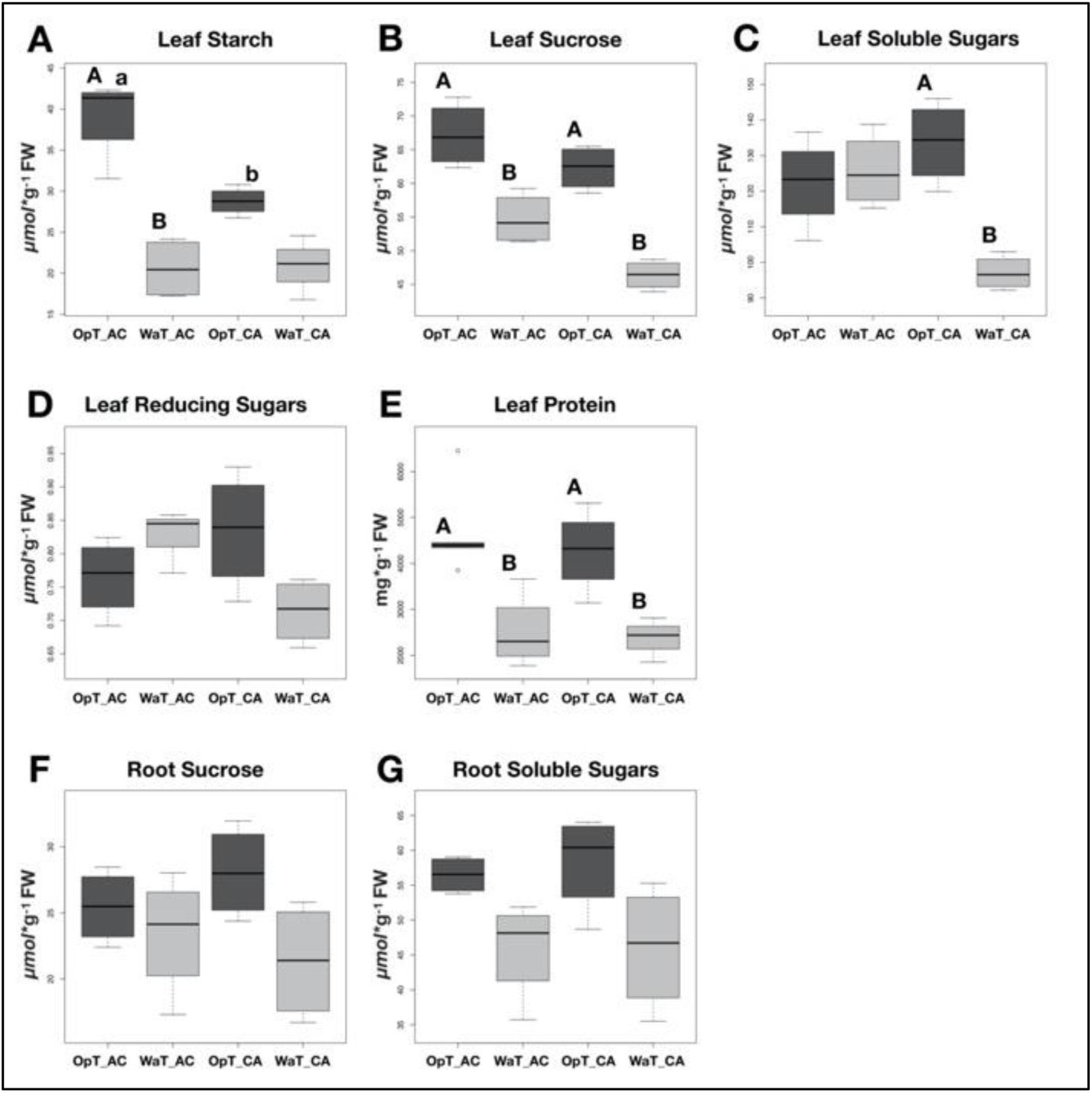
Boxplot analyses of different sugars and protein content in leaves and roots of two coffee genotypes under two temperature conditions. Coffee genotypes, cv. Acauã (AC) and Catuaí (CA) were subjected to two temperature conditions (OpT and WaT) and the starch content (A), sucrose (B), soluble sugars (C), reducing sugars (D) and protein (E) were determined in leaves. The content of sucrose (F) and soluble sugars (G) were also measured in roots. Statistical analyses were performed comparing the same coffee genotype at different temperatures (capital letters) and comparing different genotypes at the same temperature (small letters). Differences were considered significant at p<0.05 (see Table S4 for details). Labels: OpT - optimal temperature (23/19°C, day/night); WaT - warm temperature (30/26°C, day/night); CA - cv. Catuaí; AC - cv. Acauã.

For leaf sucrose content, both genotypes showed similar content at OpT conditions (Fig. 4B), but the reduction in sucrose content was significantly greater for cv. Catuaí (P<.003) when compared to cv. Acauã (P<.018). Leaf soluble sugars were only reduced in cv. Catuaí leaves under WaT conditions (Fig. 4C), whereas reducing sugar content was similar in both cultivars at both growth temperatures (Fig. 4D). Leaf protein content of cv. Acauã and cv. Catuaí mirrored one another with a marked decrease in protein content under warm stress conditions (Fig. 4E). By comparison, roots did not exhibit a difference in sucrose or soluble sugars content due to growth temperature or genotype identity (Fig. 4F and 4G).

Thus, both the carbohydrate and protein content analyses revealed that moderate increases in growth temperature impacts coffee leaves at the metabolic level. Moreover, the specific differences between coffee genotypes in response to elevated temperatures, especially starch and soluble sugars, demonstrated that temperature effect is genotype-dependent.

## DISCUSSION

Coffee genotypes differ in their physiological and molecular responses to warm temperature stress reflecting an intraspecific genetic variability that could be useful for breeding (DaMatta, 2018). To explore this variation and to better understand thermotolerance mechanisms, we compared the effect of warm temperature on the physiological, transcriptional, and metabolic status of two commercial coffee genotypes. The present study revealed that cv. Acauã showed a better thermoregulatory capacity maintaining lower temperature and transpiration foliar levels than cv. Catuaí, whereas photosynthetic rates did not differ between the coffee cultivars (Fig. 1 and Table S1). The present findings are in agreement with recent studies that describe photosynthetic stability, as well as thermoregulatory differences, between coffee genotypes in response to increased growth temperatures (Bertrand *et al*., 2015; Martins *et al*., 2016, DaMatta *et al*., 2019). Comparative studies identifying warm tolerant coffee genotypes are scarce and our results show that, based on the established experimental conditions, cv. Acauã appears to better regulate temperature than cv. Catuai for some of the physiological parameters analyzed.

The pathways that integrate temperature perception with physiological and metabolic regulation in plants largely depend on complex transcriptional networks (Ruan *et al*., 2010; Bita and Gerats, 2013). Given this, together with observed physiological differences between cv. Acauã and Catuaí, we conducted an RNAseq analyses of coffee leaves in response to elevated growth temperature. The results showed a number of DEGs (Table S1) including HSPs and genes related to photosynthesis and carbohydrate regulation, which agree with published literature from other species (Jagadish *et al*., 2016; Ohama *et al*., 2016).

One noted advantage of RNAseq analysis is the global examination of all expressed genes under defined environmental/developmental conditions. This permits a detailed examination of the entire transcriptome to reveal stimulus-driven mechanisms (Lowe *et al*., 2017). In the present study, the higher number of DEGs found between coffee genotypes at OpT compared to WaT conditions was unanticipated (Fig 2) and demonstrates that both cultivars show a similar transcriptional response to warmer temperatures. Regarding this, it is tempting to suggest a bottleneck effect of transcription in response to warm temperatures. Yu *et al*. (2007) coined the term bottleneck to refer to highly centrality regulatory nodes that play key roles in mediating communication within a given network. Here, warmer temperatures would function as a centralizing point of transcription, possibly converging similar responses for thermoregulation and relocating energy resources between pathways, thereby acting as a constraint of variability. This is in accordance with the concept of hubs (or bottlenecks) in molecular signaling networks (Dietz *et al*, 2010), and the genotype-dependent effect of ambient temperature in plant plasticity (Ibañez *et al*., 2017; Zhu *et al*. 2018). Thus, stress-imposed constraints impact the energy costs for plant development limiting phenotypic plasticity (Auld *et al*., 2010; Murren *et al*., 2015) and bringing a grand challenge to select crop genotypes more resilient to climate change (Pereira, 2016).

In response to warm temperature, we found six DEGs shared by the two coffee genotypes examined (Fig. 2D and Table S2), whose expression trends were validated by RT-qPCR (Fig. S7). These results indicate a core conservative thermoregulatory mechanism within the coffee genotypes. The homologues of *Small Heat shock Proteins* (Cc11_g16360) are triggered in response to stress and during the ripening process, acting as chaperones presenting a complex expression pattern (Ohama *et al*., 2016; Arce *et al*., 2018). *RuBisCO activase* (Cc04_g14500), whose homologues promote RuBisCO activity (Salvucci and Ogren, 1996), is up-regulated by WaT in coffee. RuBisCO is involved in CO_2_ fixation during photosynthesis and it is negatively affected by increased growth temperature (Salvucci *et al*., 2001; Crafts-Brandner and Salvucci, 2000). Thus, the increase of *RuBisCO activase* in coffee leaves was interpreted as a compensatory mechanism at WaT agreeing with photosynthetic rates that were unaffected by warmer temperature (Fig. 1C). Homologues of *Isoflavone reductase* (Cc10_g02660) are involved in isoflavonoid synthesis, which are secondary metabolites related to lignin biosynthesis and pathogen defense (Shoji *et al*., 2002; Cheng *et al*., 2015). However, the direct relationship between *Isoflavone reductase* and temperature stress has not been previously established (Wang *et al*., 2006b). The DEGs *UP-9A* (Cc02_g00580) and *TPR-like* (Cc01_g17230) appear to be a stress response related to sulfur deficiency. In *Arabidopsis*, homologues of *UP-9A* are putative interactors with ADP-glucose, which plays a key role in starch metabolism by converting glucose 1-phosphate to ADP-glucose (Crevillén *et al*., 2005). Homologues of *Granule-bound starch synthase* (Cc08_g16970) are involved in starch and sucrose metabolism pathways and in thermotolerance acquisition (Wang *et al*., 2006a; Tian *et al*., 2018).

A series of DEGs responsive to warmer temperatures were not shared between the two cultivars suggesting the possible existence of genotype-specific thermoregulatory mechanisms (Fig. 2D). We observed that many of these DEGs are involved with carbohydrates and carbon regulatory pathways (Fig. S1 to S6). Within cv. Acauã, nearly twice as many differentially expressed genes were found in WaT versus OpT in comparison to WaT versus OpT DEGs in cv. Catuaí (Fig. 2D). Of particular interest, cv. Acauã showed a downregulation of Cc10_g04070 in WaT, a gene specific to the pathway for stomatal closure via ABA regulation (Fig. S5). As plants generally close stomata to prevent excess water loss in warm temperatures, a downregulation of a gene signaling the stomata to close may indicate a reduced sensitivity to slightly warmer temperatures.

Sugar metabolism is a complex and dynamic process strongly controlled by many pathways, and once deregulated very often affects the carbon and protein partitioning in plants (Rolland *et al*., 2006; Gutiérrez *et al*., 2007; Zhang *et al*. 2012; Kölling *et al*., 2015). Because coffee genotypes differ in starch and sucrose metabolic pathways (Fig. S4), we hypothesize that, in response to warm temperatures, different enzymes would be activated triggering changes in carbohydrate and protein content. For example, cv. Acauã could accumulate more starch in leaves than cv. Catuaí under OpT conditions, which would represent a carbohydrate reserve when unfavorable growth conditions are imposed.

Our results demonstrated that there are differences in leaves regarding the sugar content, such as starch, sucrose and total soluble sugars (TSS), and in total protein content in response to warmer temperatures (Fig. 4). These negative correlations between sugar content and temperature stress are in agreement with the report for *C. arabica* L. by Bertrand *et al*. (2015), and reinforces our conclusion that warmer temperature has a genetic and physiologic impact on coffee leaves in a genotype-dependent manner.

## CONCLUDING REMARKS

Coffee is a worldwide commodity, produced in over 80 countries and foundational to the economy of many regions through employment and trade (DaMatta and Ramalho, 2006). Climatic changes will not only affect the production of this high-value commodity, it will also elicit major economic and social repercussions. In this work, we showed an overall reduction in the number of differentially expressed genes between coffee genotypes under warmer temperature in comparison to optimal temperature. Gene expression at optimal temperatures was more diverse suggesting that these genotypes have variable baseline transcription. Although our findings demonstrate potential differences in starch and sucrose metabolic pathways along with variable physiological responses among cultivars, more studies are required to substantiate patterns of coffee thermotolerance. Our results confirm the need to search and develop more climate-adaptive coffee varieties as extant *C. arabica* L. cultivars may not possess the genetic variability needed to ensure consistent and profitable production of coffee in warmer temperatures (Lashermes *et al*., 1999; Anthony *et al.*, 2001; Cubry *et al*. 2008). Further investigation on intra and interspecific elevated temperature tolerance is warranted which may be critical to inform future introgression efforts of existing stress tolerance traits from other *Coffea* species into preferred Arabica genotypes.

## SUPPLEMENTARY DATA

Primer list - Primers used for RT-qPCR.

Table S1 - Statistical analyses of physiological analyses presented in Fig. 1.

Table S2 - Differentially expressed genes and statistical analyses of RNAseq data presented in Fig. 2 and 3.

Table S3 - Statistical analyses of RT-qPCR presented in Fig. S7.

Table S4 - Statistical analyses of sugars and protein analyses presented in Fig. 4.

Fig. S1 to S6 - Functional characterization of DEGs at different conditions found by RNAseq analysis.

Fig. S7 - RT-qPCR analysis of ten selected DEGs identified by RNAseq.

## ACKNOWLEDGEMENTS

The authors thank the members of the Laboratory of Plant Molecular Physiology (LFMP, UFLA/Brazil) for structural support of the experiments; Prof. Dr. Paulo Eduardo R. Marchiori and Dra. Lissa V. Vilas Boas (UFLA/Brazil) for the support on sugars and proteins analyses; Prof. Dr. Robert R. Klein (USDA/USA) for the support on RT-qPCR analysis. This work was financially supported by the National Council of Scientific and Technological Development (CNPq) and National Institutes of Science and Technology of Coffee (INCT/Café).

## AUTHORS’ CONTRIBUTION

Project conceptualization: ACJ, BCFB, PEK.

Paper conceptualization: RRO, THCR.

Physiological analyses: RRO, THCR, CHC, CFC.

RNAseq library preparation: BCFB.

RNAseq analysis: THCR.

Bioinformatic analyses: THCR, RRO.

RT-qPCR analyses: LF, PEK.

Sugar and protein analyses: RRO, THCR, CHC.

Statistical analyses: THCR, VAM, RRO.

Writing – original draft: RRO, THCR.

Writing – review and editing: RRO, LF, PEK, ACJ.

